# The effect of the preceding masking noise on monaural and binaural release from masking

**DOI:** 10.1101/2021.11.05.467091

**Authors:** Hyojin Kim, Viktorija Ratkute, Bastian Epp

**Affiliations:** Hearing Systems Section, Department of Health Technology, Technical University of Denmark, Kgs. Lyngby, Denmark; Biomedical Engineering, Technical University of Denmark, Kgs. Lyngby, DK-2800, Denmark

## Abstract

Auditory stream segregation can be facilitated when the maskers share coherent amplitude modulations or by utilizing spatial cues. The effectiveness of each cue can be quantified as a decrease in masked thresholds, termed as comodulated masking release (CMR) and binaural masking level difference (BMLD). Prolonged exposure to the masker can influence following target segregation. However, the collective impact preceding noise on target segregation in the presence of comodulation and interaural phase difference (IPD) cues is unclear. Stimuli were used to induce noise streams by altering the duration and temporal coherence of the envelope of the preceding masker. The effect on following target detection with CMR and BMLD induced by an IPD of the target tone was investigated. The results indicate that the effect of the preceding stream formation on CMR operates on different time scales, extending beyond 200 ms, depending on the spectrotemporal characteristics of the maskers. However, the effect on IPD-induced BMLD was not significant across time. Under the simplifying assumption that peripheral processing operates on shorter time scales than cortical processing, the results of the present study may provide insights into auditory signal processing in the presence of beneficial cues for target stream segregation.

## 1. Introduction

The ability to listen to a particular talker in a noisy environment is one of the most outstanding abilities of the auditory system of humans. This ability is often referred to as the “cocktail party effect” [Cherry, 1953]. This effect can be interpreted as “auditory scene analysis”, suggesting that our auditory system perceives the acoustic environment as a combination of multiple auditory streams [Bregman, 1994]. By combining various sound features extracted from the sounds reaching the two ears, the auditory system can form auditory streams, enabling us to segregate a target stream from a noisy background. This is analogous to the visual system that uses visual cues such as colors, textures, and shapes to separate a target object from other visual objects. Hence, a stream can be thought of as a basic unit of sound perception resulting from a perceptual grouping of auditory cues [Bregman, 1994]. The question is how various auditory cues are combined along the auditory pathway and play a role in inducing perceptual auditory streams collectively.

Based on psychoacoustical studies, one can speculate that sound elements that come from the same sound source can be a grouping cue to form a auditory stream [Moore and Gockel, 2012]. Firstly, all frequency components emitted from the vibration of the vocal tract during speech production share the same temporal modulation pattern [Raphael et al., 2007]. This shared temporal modulation is referred to as comodulation and this is a common property of natural sounds, suggesting that comodulation may play a role as a grouping cue for the auditory system [Nelken et al., 1999]. The benefit of comodulation has been shown experimentally with a target detection task in noise [Hall et al., 1984, 1990]. The masked threshold of a target tone decreases when a masking noise is comodulated compared to the condition where the target tone is masked by the noise with an uncorrelated modulation pattern. This enhancement in the target detection performance is commonly referred to as comodulation masking release (CMR) [Hall et al., 1984]. In the light of stream formation, it can be argued that comodulation supports the grouping of frequency components of the masker into one stream, thereby facilitating the segregation of the target tone from the masker.

Secondly, sound components coming from the same position in space share spatial information, which can also facilitate following a particular speaker in a noisy environment [Bregman, 1994]. Similar to comodulation across frequency, spatial information in the form of interaural differences can induce a masking release. When sounds reach the ears from a specific location in space, differences in phase and level between the ears are introduced. These interaural differences can be beneficial for target detection. This can result in a decrease in masked thresholds induced by interaural disparities, which can be quantified as the binaural masking level difference (BMLD) [Jeffress et al., 1956, van de Par and Kohlrausch, 1999].

In physiological studies, neural correlates of masking release are often reflected as enhancement or suppression of neural signals. For CMR, neural correlates have been found at various stages along the auditory pathway. The earliest neuronal encoding of CMR was found in the cochlear nucleus (CN) as a reduced responses to the amplitude-modulated masker compared to uncorrelated noise [Pressnitzer et al., 2001]. As a possible neural mechanism, they suggested an across-channel processing where a wideband inhibitor inhibits the responses to the amplitude-modulated masker while not affecting the responses to the target signal. This results in an increased spike rate change in response to the target signal between the masker-only and masked signal conditions when the masker is comodulated compared to the condition with uncorrelated masker [Pressnitzer et al., 2001, Neuert et al., 2004]. Other neural correlates were found along various stages along the axis of the midbrain to the auditory cortex (A1). Along this axis, the neural representation of the target tone in modulated noise shows gradual changes, suggesting the emergence of neural representations of auditory objects: Neural units in the inferior colliculus (IC) show both a reduced response to the masker and an enhanced response to the target tone when the masker is comodulated [Diepenbrock et al., 2017]. In addition, from the medial geniculate body (MGB), a new neural representation arises as an envelope locking suppression (ELS) where neuronal locking to the envelope of the modulated noise is suppressed with an addition of the target tone [Diepenbrock et al., 2017, Las et al., 2005, Nelken et al., 1999]. Overall, these findings indicate that along the IC-MGB-A1 axis, the neural representations of auditory objects arise in the presence of fluctuating noise. For BMLD, physiological correlates of binaural processing were found at in superior olivary complex (SOC) [Grothe et al., 2010, for a review], the dorsal nucleus of the lateral lemniscus (DNLL) [e.g. Siveke et al., 2006], and the IC where input from both ears strongly converges [Shackleton et al., 2005, 2003, Zohar et al., 2011].

A previous psychoacoustic study showed the superposition of CMR and BMLD where the overall masking release introduced by comodulation and IPD was close to the sum of CMR and BMLD [Epp and Verhey, 2009b]. This was interpreted and modeled as a serial improvement of the masked signal along the CN-IC axis. However, neither the exact order of CMR processing and IPD processing nor the physiological basis underlying these effects could be fully identified. In addition, in some cases, listeners with high BMLD showed a reduction in the amount of CMR [Epp and Verhey, 2009a, Hall III et al., 2011, Egger et al., 2019]. Thus, it remains unclear whether the neural representations of comodulation and IPD cues are linearly combined along the CN-IC axis. Previous studies have not considered interaural disparities in the masker, which could have different implications. Moreover, it is unclear how neural enhancements associated with CMR and BMLD contribute to the target segregation along the IC-MGB-A1 axis.

Besides CMR and BMLD, target detection performance can also be affected by a “build-up” period where the auditory system can collect sound information over time. This phenomenon is referred to in various terms such as “build-up of stream segregation”, “adaptation to auditory streaming”, or “temporal decline of masking” [Moore and Gockel, 2012, Anstis and Saida, 1985, McFadden and Wright, 1992]. In combination with CMR, several studies have shown that presentation of a precursor can decrease or increase the amount of CMR, depending on the spectrotemporal characteristics of preceding maskers [Grose et al., 2009, Dau et al., 2005]. Physiologically, neural correlates of the influence of preceding maskers on CMR have been found in the primary auditory cortex (A1) [Sollini and Chadderton, 2016]. While this study supports the idea that the effect of precursors originates from the cortex level, they also suggested the role of A1 in cortical feedback to subcortical regions, affecting the bottom-up processing of the preceding maskers. For BMLD, physiological evidence indicates the existence of IPD neural processing at IC level, however, little is known if the presence of preceding maskers, or stream formation, can affect following spatial hearing later in time. Based on these findings, one can speculate that binaural processing plays an important role in stream formation by spatial disparities. However, binaural processing can be precise in encoding small interaural differences while it can be imprecise in processing over longer time scales. These include motion aftereffects [Grantham and Wightman, 1979] and slow processing of changes in interaural cues, often referred to as “binaural sluggishness” [Kollmeier and Gilkey, 1990, Eurich and Dietz, 2023]. Furthermore, in the light of the phenomena linked to informational masking, it is unclear how binaural processing explicitly contributes to the stream formation [Middlebrooks and Simon, 2017, for an overview].

Thus, we conducted further investigations into the effect of the preceding stream formation on both CMR and IPD-induced BMLD under two simplifying assumptions. The first assumption was that the neural representation of the auditory stream arises from the IC-MGB-A1 axis on top of the encoding of comodulation and IPD occur along the CN-IC axis. The second assumption was that neural processing operates on increasingly longer time scales as the neural signal propagates from the periphery (short time constants) to cortical areas (long time constants). Based on these assumptions, we hypothesized that:

i. If the neural encoding of stream formation is serially added to the encoding of CMR and IPD-induced BMLD along the IC-MGB-A1 axis, the preceding stream formation will affect both CMR and BMLD.
ii. If the stream formation results from higher-order processing, the effect of the preceding stream formation on the following target detection will gradually increase with longer priming periods to the preceding maskers, exhibiting a relatively long time constant.

To test these hypotheses, we designed four masker types to induce various preceding stream formation where each condition provides grouping cues between maskers inducing one or multiple streams. We varied the duration of the preceding maskers to determine the time scale of the interaction of preceding stream formation with CMR and IPD-induced BMLD.

## 2. Methods

### 2.1. Stimuli

The stimulus consisted of a preceding masker interval with varying duration and a subsequent masked tone interval lasting 200 ms including 20 ms raised-cosine on- and offset ramps (Fig. 1). To avoid audible artifacts, the preceding masker interval was overlapped with the masker with the target tone interval using a 20 ms raised-cosine off- and onset ramp with 50% overlap. The preceding masker interval duration were 0, 20, 100, 200, 300, or 500 ms including 20 ms raised-cosine onset ramps. Each masker consisted of five narrow-band noises with 20 Hz bandwidth. The center band (CB) was centered at 700 Hz and the remaining four masker bands (flanking bands, FBs) were centered at 460, 580, 820, and 940 Hz. The spectral distance between the masker bands was chosen to maximize CMR based on Grose et al. [2009]. The overall level of the masker was set to 60 dB SPL. The masker was in all conditions presented identically to both ears.

**Figure 1:**
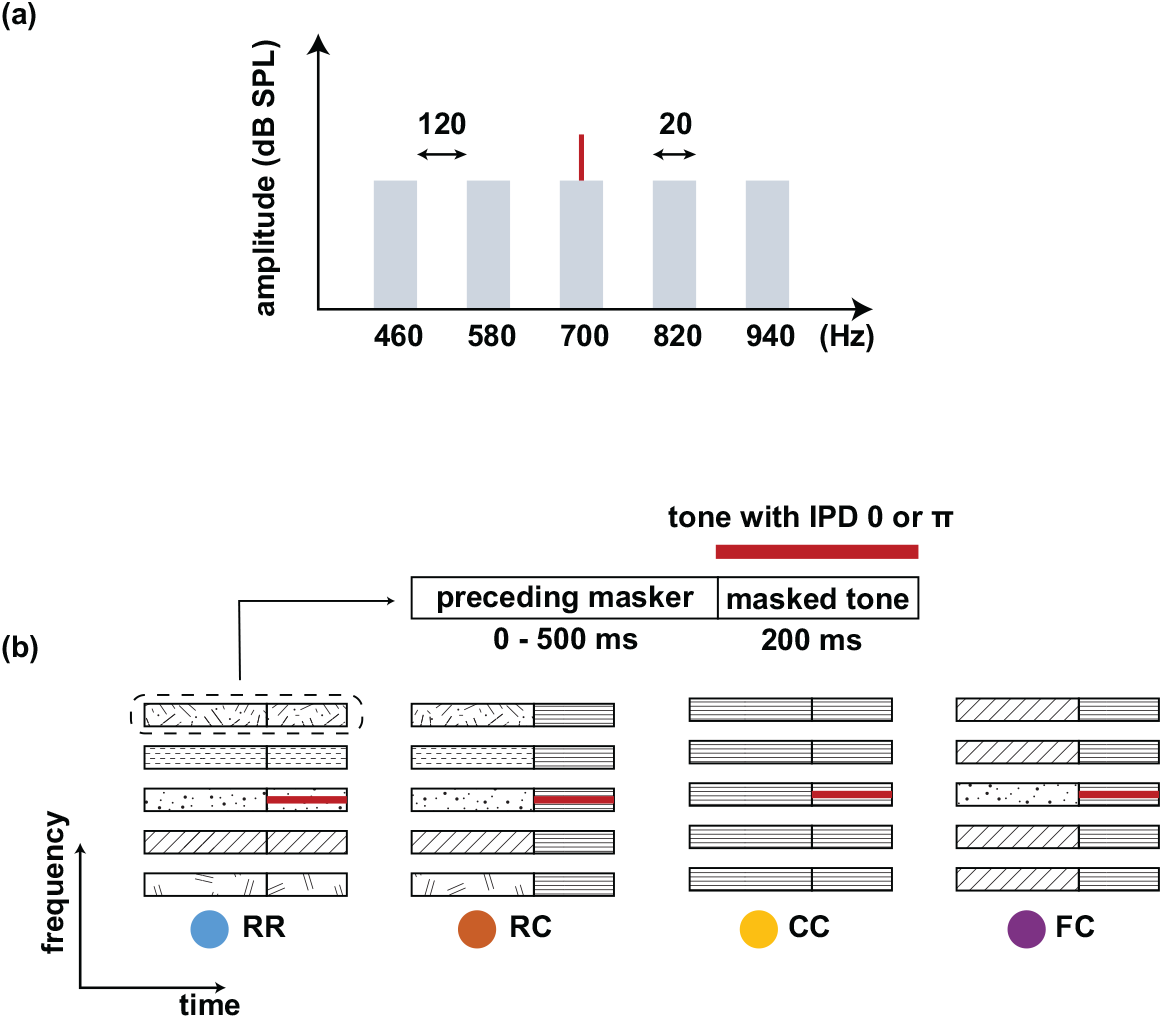
(a) Spectra of the stimulus. A target tone (700 Hz) was presented with a masking noise consisting of five narrow-band maskers: One signal centered band (SCB) and four flanking bands (FBs). The bandwidth of each masker band was 20 Hz, and the frequency spacing was 120 Hz. The overall level of the noise was set to 60 dB SPL. (b) Schematic spectrograms of the stimulus conditions. Each stimulus consisted of a preceding masker interval (0 - 500 ms) and a masked tone interval (200 ms). Four types of maskers were used: RR, RC, CC, and FC. The RR was used as the reference condition with uncorrelated masker bands. In the other three conditions, the maskers consisted of a comodulated masker preceded by three different maskers: uncorrelated masker (RC), comodulated masker (CC), and the masker with comodulated flanking-bands (FC). The thick line represents a tone that was presented with an IPD of 0 or *π*.

Firstly, we designed two conditions with no preceding masker (0 ms) prior to the target tone. In these two conditions, the masker had either uncorrelated intensity fluctuations across frequency (R), or coherent intensity fluctuations across frequency (comodulated, C). Secondly, to investigate the effect of preceding stream formation on following masking release by comodulation and IPD cue, we used four different masker types: i) uncorrelated masker for both the preceding and following maskers (RR). The three remaining conditions consisted of a comodulated masker (C) preceded by ii) an uncorrelated masker (RC), iii) a comodulated masker (CC), or iv) a masker with comodulated flanking-bands (FC). The preceding R masker bands only contained common onset as a cue. Hence, we used the RR masker to estimate the effect of preceding masker energy (here referred to as adaptation effect) on the following target detection interval. For the RC masker, we hypothesized that the preceding uncorrelated masker bands only induce the adaptation effect on the following comodulation cue for the target segregation from the masker. This will allow to shed light on the interplay between the adaptation effect and CMR in the masked tone interval. For the CC masker, the preceding masker bands would be grouped together into one masker stream and facilitate following masking release, as in a classical CMR flanking band paradigm. For the FC masker, only the FBs would be grouped into one stream, hindering following masking release as the comodulation in the masked target interval would conflict with the preceding masker. To test the interaction of between CMR and IPD-induced BMLD, the target tone was presented with an interaural phase difference (IPD) of 0 (diotic conditions) or *π* (dichotic conditions) in combination with the four masker types presented diotically. Here, for instance, *CC*_0_ indicate a condition with the CC masker type and IPD of 0, and *CC*_*π*_ indicate a condition with the CC masker type and IPD of *π*.

### 2.2. Protocol

All listeners performed one training session which included all conditions. Then, each listener performed three threshold measurements for all conditions. For each measurement, all eight conditions were presented in randomized order. The thresholds were estimated by averaging the three threshold measurements for each condition. An additional measurement was done if the thresholds from the three measurements had high variance (SD > 3 dB). During the threshold measurement, the listeners were seated in a double-walled, soundproof booth with ER-2 in-ear headphones. We used an adaptive, three-interval, three-alternative forced-choice procedure (3-AFC) with a one-up, two-down rule to estimate the 70.7% of the psychometric function [Ewert, 2013, Levitt, 1971]. Three sound intervals were presented with a pause of 500 ms in between. Two intervals contained only the maskers, while one interval contained the target tone with maskers. The listeners’ task was to choose the interval with a target tone by pressing the corresponding number key (1, 2, 3) on the keyboard. Whenever the listener pressed the keyboard, visual feedback was provided, indicating whether the answer was “WRONG” or “CORRECT”. The target tone’s start level was set to 75 dB. Depending on the answer, the target tone level was adjusted with an initial step size of 8 dB. The step size was halved after each lower reversal until it reached the minimum step size of 1 dB. The signal level at a minimum step size of 1dB was measured six times, and the mean of those last six reversals was used as the estimated threshold.

### 2.3. Listeners

Ten normal-hearing listeners participated in the experiment (5 male, 5 female). All participants were under the age of 30 (mean: 27.7). All participants were screened with audiological hearing screening including visual inspection of the ear canal and standard audiometry. None of them reported any history of hearing impairment and had pure-tone hearing thresholds within 15 dB HL for the standard audiometric frequencies from 125 to 4000 Hz. All participants provided informed written consent, and all experiments were approved by the Science-Ethics Committee for the Capital Region of Denmark (reference H-16036391).

## 3. Results

### 3.1. Masked thresholds

We measured masked thresholds of a target tone in the presence and absence of an IPD in conditions with various masker types. Figure 2 (a-c, left panels) shows the mean thresholds across all listeners. Masked thresholds in diotic conditions (IPD of 0) are plotted with solid lines. Masked thresholds in dichotic conditions (IPD of *π*) are plotted with dotted lines. Each panel shows the masked thresholds as function of the duration of the preceding masker (20 ms to 500 ms) for four masker types: *RR, RC, CC*, and *FC*. The two masker types R and C without preceding maskers (0 ms) were used as the reference to estimate the effect of preceding stream formation on following target detection. Mean thresholds for a diotic tone and no preceding maskers were 52.8 dB SPL (*R*_0_) and 45.7 dB SPL (*C*_0_). The introduction of an IPD reduced the masked threshold by 11 to 15 dB, resulting in thresholds of 37.4 dB SPL (*R*_*π*_), and 33.6 dB SPL (*C*_*π*_). The addition of short preceding maskers (20 ms) increased the thresholds by 2.3 dB and 2.5 dB for *RR*_0_ and *RR*_*π*_ conditions, respectively (Fig. 2a). Only *RR*_0_ showed a significant difference (two-sample t-test, p < 0.05) between the absence and the presence of the preceding masker. Preceding maskers longer than 20 ms did not show a significantly increase of the masked thresholds.

**Figure 2:**
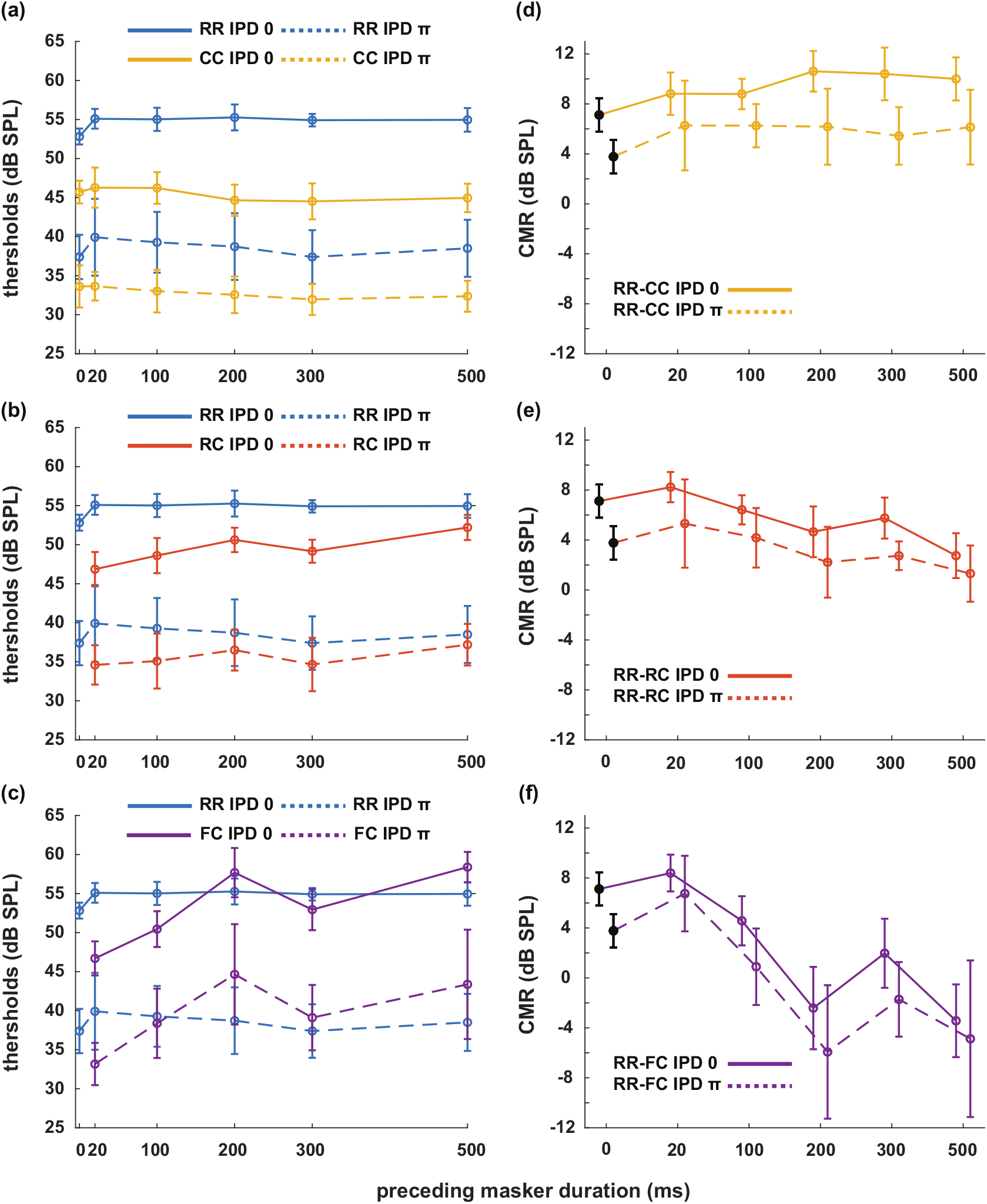
Mean masked thresholds (left column) and CMR values (right column) for each masker condition with the RR masker as a reference. Data from all listeners were averaged. Data are plotted for each masker type with colors (RR - blue, RC - orange, CC - yellow, FC - purple). For diotic conditions with an IPD of 0, masked thresholds are plotted with solid lines. For dichotic conditions with IPD of *π*, masked thresholds are plotted with dotted lines. Error bars indicate plus-minus one standard deviation.

The addition of 20 ms of different preceding masker types to a comodulated masker in the masked tone interval (*RC, CC, FC*) showed no significant changes in the masked thresholds relative to the conditions with no preceding maskers. For *CC*_0_ and *CC*_*π*_ conditions, longer preceding maskers had no significant effect on the masked thresholds for both diotic and dichotic conditions (Fig. 2a). For the *RC*_0_ condition, masked thresholds increased significantly when the preceding masker duration was longer than 200 ms (two-sample t-test, p < 0.01), reaching 52 dB at the duration of 500 ms (Fig. 2b). For the *RC*_*π*_ condition, the mean threshold reached 37.2 dB SPL when the preceding masker duration was 500 ms, showing a significant increase relative to a 20 ms preceding masker (two-sample t-test, p = 0.0386). For *FC*_0_ and *FC*_*π*_, the thresholds increased significantly as well when the preceding masker was longer than 100 ms (two-sample t-test, p<0.05). For *FC*_0_ and *FC*_*π*_ thresholds were 58.4 dB SPL and 43.4 dB SPL in combination with a 500 ms preceding masker.

We examined three factors on the masked thresholds: comodulation, IPD, and the duration of the preceding maskers. The analysis revealed a significant effect of the comodulation factor (three-way ANOVA, *F* = 392.15, p< 0.001), a significant effect of the IPD factor (three-way ANOVA, *F* = 2273.67, p< 0.001), and a significant effect of the factor duration of the preceding maskers (three-way ANOVA, *F* = 17.81, p< 0.001). There was a significant interaction between the comodulation IPD factors (three-way ANOVA, *F* = 8.27, p<0.001). There was also a significant interaction between the comodulation and duration (three-way ANOVA, *F* = 13.23, p< 0.001). However, there was no significant interaction between the IPD and the duration of the preceding maskers.

### 3.2. Comodulation masking release

Using the above thresholds measurements, we studied the effect of preceding stream formation on CMR while manipulating the duration of the preceding masker. We calculated CMR for three conditions (*CC, RC*, and *FC*) by subtracting the corresponding thresholds from the thresholds of the reference conditions (*RR*). Here, corresponding CMR measures are denoted as *CMR*_*CC*_, *CMR*_*RC*_, and *CMR*_*RC*_, respectively. Figure 2 (right panels, d-f) shows changes in CMR as a function of the duration of preceding maskers. As a reference, we calculated CMR for the condition with no preceding masker by subtracting the masked threshold in the *C* condition from the threshold in the *R* condition (*CMR*_*C*_). CMR in the diotic condition (*CMR*_*C*0_) was 7.1 dB while CMR in the dichotic condition (*CMR*_*Cπ*_)was 3.8 dB. For a short preceding masker (20 ms), all conditions showed a slight increase in CMR of around 1.5 dB in diotic conditions and of 2.4 dB in dichotic conditions relative to the absence of the preceding masker.

We tested the significance of the changes in CMR by changing the duration of the preceding masker from 20 ms to 500 ms. Both *CMR*_*CC*0_ and *CMR*_*CCπ*_ did not show a significant change. *CMR*_*RC*0_ and *CMR*_*RCπ*_ both showed a significant decrease from 8.23 dB (20 ms) to 2.75 dB (500 ms) (one-way ANOVA: *F* = 16.17, *p* < 0.01), and from 5.32 dB (20 ms) to 1.31 dB (500 ms) (one-way ANOVA: *F* = 3.9, *p* = 0.008), respectively. A multiple comparison test with Bonferroni correction showed that *CMR*_*RC*0_ significantly reduced when the duration of the preceding maskers was longer than 200 ms (p < 0.05). *CMR*_*RCπ*_ significantly reduced when the duration of the preceding maskers was 500 ms (p < 0.05). *CMR*_*FC*0_ showed a significant decrease from 8.39 dB (20 ms) to -3.43 dB (500 ms) (one-way ANOVA: *F* = 36.47, *p* < 0.01), and *CMR*_*FCπ*_ showed a significant decrease from 6.75 dB (20 ms) to -4.87 dB (500 ms) (*F* = 13.57, *p* < 0.01). A multiple comparison test with Bonferroni correction showed that *CMR*_*FC*0_ significantly reduced when the duration of the preceding maskers was longer than 100 ms (p < 0.01). *CMR*_*FCπ*_ significantly reduced when the duration of the preceding maskers was longer than 100 ms (p < 0.05).

### 3.3. Binaural masking level difference

Lastly, we investigated the effect of a preceding stream formation on BMLD. We calculated BMLD for four conditions (*RR, RC, CC*, and *FC*) by subtracting the masked threshold of dichotic conditions from those of the corresponding diotic conditions (Fig. 3). Here, corresponding BMLD measures are denoted as *BMLD*_*CC*_, *BMLD*_*RC*_, and *BMLD*_*RC*_, respectively. As a reference, we calculated BMLD for the condition with no preceding masker by subtracting the masked threshold of the dichotic condition from the threshold of the diotic condition. BMLD for a random masker (*BMLD*_*R*_) was 15.4 dB and for a comodulated masker (*BMLD*_*C*_) 12.1 dB. BMLD did not show a significance change (one-way ANOVA) with varying the duration of the preceding maskers. Additionally, *BMLD*_*FC*_ exhibited higher variance compared to other masker types when the duration of the preceding maskers were 200*ms*, and 500*ms* (*F − test*, p<0.05). We observed that the standard deviation of BMLD measures increased from 2 dB to 6.7 dB, while the standard deviations of the other conditions were between 2 and 3.8 dB.

**Figure 3:**
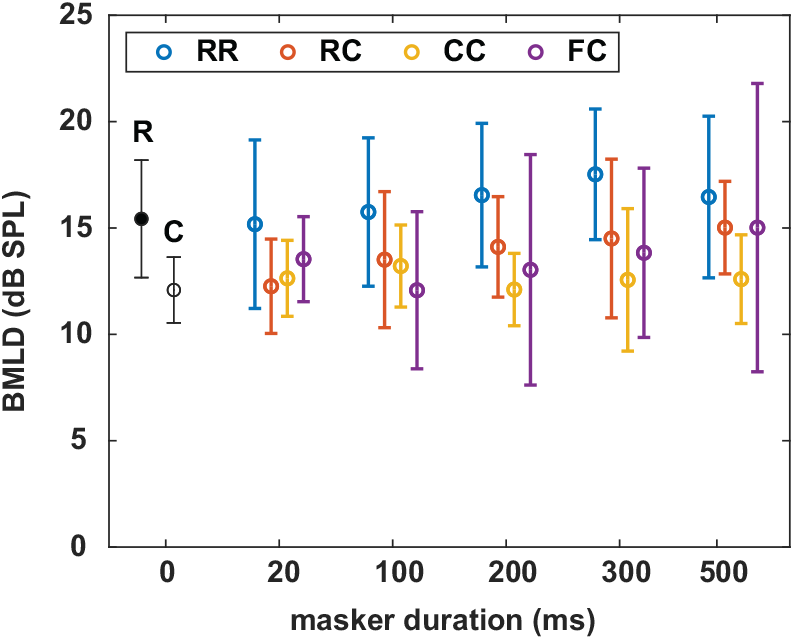
Mean BMLD for each masker condition. Data from all listeners were averaged. Data are plotted for each masker type with colors (RR - blue, RC - orange, CC - yellow, FC - purple). As a reference, BMLD in the R and C conditions are shown in black.

## 4. Discussion

We investigated the effect of preceding stream formation on following target detection. We applied the simplifying assumption that physiological processing shows longer time constants for more central relative to more peripheral processing stages. We used CMR and IPD-induced BMLD as measures of the target detection performance. We designed four types of the preceding maskers to induce various streaming conditions. We used different combinations of maskers with and without comodulation as a grouping cue in the preceding masker interval and the masked tone interval (*RR, CC, RC*, and *FC*). In the *RR* and *RC* conditions, only common onset, but no comodulation was available in the preceding masker interval to form a stream. In the *CC* and the *FC* conditions, comodulation provided an additional cue to stream formation for either all masker bands (*CC*) or the flanking bands only (*FC*). We varied the duration of the preceding maskers, and estimated the effect of the preceding stream formation on the following release from masking of the target tone, facilitated by comodulation(CMR) and interaural phase disparity of the target tone (IPD-induced BMLD). Our hypotheses were, that if a stream formation is the result of the high-level processing along the IC-MGB-A1 axis, CMR and BMLD measures will be affected by the preceding maskers, and the effect size will increase with longer priming periods to the preceding maskers.

### 4.1. The effect of preceding stream formation in diotic conditions

If a stream formation is the result of high-level processing, the effect of preceding stream formation on masking release will increase by longer duration of the preceding maskers as prolonged exposure to the maskers will provide more information to the central processing. For *CMR*_*RC*0_, data show that the preceding noise can disrupt the benefit of the following comodulation cue (> 200 ms). This can be interpreted as the result of adaptation effect by preceding random noise on the following target detection. However, *RR*_0_ conditions only showed minimal increase in thresholds around 2 dB even with the longest preceding maskers, excluding the adaptation effect by preceding random noise. With an assumption that uncorrelated masker does not induce a stream, excluding the possibility of the high-level processing, a reduction in *CMR*_*RC*0_ reaching up to 5.5 dB at 500 ms may indicate a negative effect on the activation of a wideband inhibitor at the CN level. For *CMR*_*CC*0_, we assumed that the preceding maskers will form one stream of noise, facilitated by both common onset and by comodulation. This would facilitate the following target detection. However, there was no significant improvement in the amount of CMR for prolonged exposure to the preceding maskers from 20 ms to 500 ms. This indicates that common onset and comodulation were sufficient and the preceding stream formation did not provide any additional benefit.

For *CMR*_*FC*0_, data show that the preceding stream formation of FBs has strong impact with the comodulation cue in the masked tone interval. This might indicate that the preceding stream formation can disrupt the following target segregation from the noise. *CMR*_*FC*0_ is reduced by up to 11.8 dB for a duration of the preceding masker of 500 ms. With an assumption that comodulated FBs form a separate stream from the centered band, relatively longer time span up to 500 ms and the strong reduction of CMR compared to other conditions may indicate an influence of high-order processing. Assuming that the effect of pre- and postcursors is the same, this finding is consistent with e.g. [Dau et al., 2009]. While the argument is strong for postcursors (due to the fact that the postcursor is presented after the target), the contribution of peripheral processing can only be ruled out if potential top-down effects to peripheral stages are excluded with a time constant of 500 ms. Some listeners also reported that they could hear the target tone for all three sound stimuli when only one stimulus contains the target tone during the listening task. As the centered noise band has the width of 20 Hz, this may have induced a tonal-like perception. Based on these results, we suggests that a stream formation at high-level processing must result in “perceptual stream”. For instance, in case of the *CC* masker, the preceding stream formation with comodulation may only result in reduced masker level perception [Verhey and Heeren, 2015], but not necessarily form a perceivable stream. If that is the case, the interaction of preceding stream formation and CMR might not increase with prolonged exposure to the preceding masker duration. Similarly, the *RC* masker does not induce a preceding stream, and since no adaptation effect of the random noise on the following target tone was observed from *RR* conditions, a decrease in *CMR*_*RC*_ may be attributed to the inhibiting mechanisms on the activation of a wideband inhibitor at the CN level. While the existence of the inhibition mechanism on the CMR processing at the CN level has not been tested, this can be supported by the physiological evidence showing the multisecond-adaptation at the CN to tonal stimuli [Pressnitzer et al., 2008]. If this is the case, our results may indicate that even the low-level processing can show a longer time constants (> 200ms) even if it does not result in perceptual streams.

McFadden and Wright [1992] measured masked thresholds with stimuli consisting of preceding maskers and a following target tone of 25 ms or 240 ms duration. This study also varied the duration of the preceding maskers and compared masked thresholds between the correlated and uncorrelated conditions, which are comparable to *CC* and *FC* conditions in the present study, respectively. In their study, the difference in a “temporal decline of masking” between two conditions reached up to 10 dB when the longest preceding masker (400 ms) was used. While the difference between two conditions is comparable to the present study, both conditions showed a “temporal decline of masking” of 4 - 6 dB in the uncorrelated condition and 15 dB in the correlated conditions. On the contrary, in the present study, *CC* conditions did not show any significant changes and *FC* conditions showed a significant reduction in masking release about 12 dB. In their study, the difference in the stimuli from the present study was primarily in the bandwidth of the noise band (50 Hz) and the target signal frequency (1250 Hz) [McFadden and Wright, 1992]. We speculate that the uncorrelated conditions in their study might not induce a tonal-like perception due to their wider noise bandwidth.

### 4.2. The effect of preceding stream formation in dichotic conditions

In dichotic conditions, the magnitude of CMR decreased compared to the diotic conditions. For each masker type (*CC, RC, FC*), we pooled CMR measures across all preceding masker duration from 20 ms to 500 ms. Then we tested the significance in the difference between CMR measures in diotic and dichotic conditions for each masker type. All masker types showed significant differences in the median (Wilcoxon signed rank test, p<0.001). Median values of CMR were *CMR*_*CC*0_ = 9.6 dB, *CMR*_*CCπ*_ = 5.7 dB (reduction of 3.9 dB), *CMR*_*RC*0_ = 5.8 dB, *CMR*_*RCπ*_ = 3 dB (reduction of 2.8 dB), *CMR*_*FC*0_ = 1.4 dB, *CMR*_*FCπ*_ = -1 dB (reduction of 2.4 dB). Therefore, there was a consistent decrease in CMR when the target tone was presented dichotically compared to when it was presented diotically. This result supports the idea that the summation of the encoding of comodulation and IPD may not be linear, aligning with the findings of Hall III et al. [2011]. It is important to note that, given the paradigm in this study, no distinction can be made regarding potential differences between within- and across-channel CMR. *CMR*_*CCπ*_ did not show a significant decrease with an increase of the duration of the preceding maskers. *CMR*_*RCπ*_ showed a significant decrease of up to 4 dB when increasing the preceding masker from 20 ms to 500 ms. For *CMR*_*FCπ*_, the reduction in CMR was 11.6 dB, similar to *CMR*_*FC*0_ at 500 ms. This observation aligns with the hypothesis, based on the simplifying assumptions, that the decline in *CMR*_*FC*_ arises from a higher-level processing due to its interaction effects at time scales of hundreds of milliseconds.

Interestingly, in contrast to our hypothesis that high-level processing would impact both CMR and IPD-induced BMLD, we observed no difference in IPD-induced BMLD with varying the duration of the preceding maskers. This finding suggests that the temporal context of diotic presentation does not influence dichotic detection, consistent with previous studies by Epp and Verhey [2009b] and Buss and III [2011]. Notably, *BMLD*_*FC*_ exhibited higher variance compared to other masker types with prolonged exposure to preceding maskers. One possible explanation is that higher-level processing affects comodulation processing as a top-town influence, but not IPD processing. With the *FC* masker, where the center band induces a tonal-like perception, the listeners actively utilizing the binaural cue (“where”) may outperform those relying on the frequency cue (“what”). If this is the case, it could be speculated that the IPD cue for the target tone may not be affected by the preceding noise but rather by the attention, and that the interaction might be different if interaural disparities where imposed on the noise rather than the target tone. However, as our current study design does not allow to further look into these ideas. Under the assumption that preceding maskers and the comodulation and IPD cues provide conflicting information for stream formation, the task of detecting the target tone might become challenging for the listener. This might explain the increased variability in the BMLD (see Fig. 3). And while no strong learning effects have been observed during data collection, extensive training might have an impact on both the variability of the data and the overall thresholds.

### 4.3. Effect of masker configuration on CMR

The effect of spectro-temporal configuration of masker and target tone on CMR was previously shown by e.g. Hatch et al. [1995] and Grose et al. [2005]. Hatch et al. [1995] found that CMR, calculated as the difference in threshold between the absence and presence of a comodulated masker, was larger for a continuously presented masker compared to a gated masker. For a comparable stimulus configuration with four flanking bands (their five-band condition), the difference between continuously presented and gated masker was about 6 dB on average. The effect of masker duration on CMR in the CC condition in the present study was around 3 dB. The target signal frequency in Hatch et al. [1995] was 2000 Hz with a duration of 400 ms, while it was 700 Hz with a duration of 200 ms in the present study. For these frequencies, a slightly smaller CMR could be expected for the 700 Hz tone compared to the 2000 Hz tone [Schooneveldt and Moore, 1987, for a single masker band a monotic presentation]. The bandwidth and spacing of the masker bands relative to the target signal frequency was comparable. The difference in the effect of the preceding masker could be due to the short duration of the target signal in the present study and the relative to a continuous masker, short maximum duration of the preceding masker of 500 ms. Another factor that could underlie differences in thresholds is that the masker continued after the offset of the target signal, while the masker ended simultaneously with the target signal in the present study. Grose et al. [2005] investigated the effect of a temporal fringe on CMR for single- or multi-tone target signals. In their study, the temporal fringe consisted of narrow band maskers with uncorrelated envelopes (similar to the RR condition in the present study) before and after the interval with the target signal. Their data show that CMR is highly reduced or even abolished in the presence of the fringe. The reduction of CMR (RR-RC) in the present study is smaller compared to Grose et al. [2005]. This could be due to the absence of a temporal fringe after signal offset and the larger CMR in the condition without temporal fringe in the present study.

Another factor shown to influence CMR in the presence of pre-cursors is temporal coherence between the masker components. Christiansen and Oxenham [2014] presented periodically gated narrow band masking noises as pre-cursors. They found a release from masking when the masker envelopes were comodulated compared to when they showed uncorrelated envelope fluctuations. When shifting the relative timing of the on frequency masker component and the flanking band maskers, the masking release was reduced. They argued that the potential grouping of all masker bands by comodulation is disrupted by temporal incoherence of the gating of the masker bands. This finding could be remotely connected to the CMR RR-FC in the present study which, consistent with the results from Christiansen and Oxenham [2014], is reduced in the presence of a preceding masker with 500 ms duration. While consistent in the reduction of a masking release, the results might be hard to compare directly because comodulation and the temporal gating used in Christiansen and Oxenham [2014] are two competing cues for stream formation. Even when considering gating as type of envelope modulation cue, it will introduce additional spectral cues due to the sharp transition which are absent in the present study and the study by Grose et al. [2009] which might explain differences in the size of CMR between the studies.

Our main hypothesis was that if the preceding stream formation is high-level processing along the IC-MGB-A1 axis, the following target detection will be affected by prolonged exposure to the preceding maskers. If the higher-level processing occurs after the summation of comodulation and IPD encoding along the CN-IC axis, both CMR and BMLD would be affected by the preceding stream formation. As IPD-induced BMLD was hardly affected with changes in the duration of preceding maskers, our results support the notion of interaction between subcortical processes (e.g., the CN) and cortical neural processes at the A1 level [Pressnitzer et al., 2008]. It could be also argued that BMLD induced by interaural disparities in the masker could have distinct implications. In this scenario, the binaural system would need to decode the masker information when interaural masker information changes. This phenomenon might be linked to effects observed on longer time scales known to be present in the binaural system, such as binaural sluggishness Eurich and Dietz [2023]. On a neural level, Epp et al. [2013] and Egger et al. [2019] found in a combined CMR and BMLD paradigm, that the N1 component of late auditory evoked potentials (LAEPs) reflected binaural processing, whereas only the later P2 component reflected the effect of masker comodulation. Based solely on timing of LAEPs, together with an assumption that LAEPs reflect internal signal-to-noise representation of the masked signal, these findings would suggest that comodulation processing occurs after binaural processing. Additionally, using similar stimuli as the present study, a recent study by Kim and Epp [2023] showed a correlation between perceived signal-to-noise (pSNR) and the P2 component of LAEPs. Notably, for the *FC* dichotic condition, the P2 component was found to be more reliable than the N1 component as a neural measure of pSNR. This finding supports our interpretation that the reduced CMR observed in *FC* conditions with prolonged exposure to the preceding masker is the outcome of higher-order processing. While employing the same paradigm with varying the duration of preceding maskers for LAEP measures could provide additional insights into the underlying neural mechanisms of preceding stream formation effects, measuring LAEPs elicited by the subsequent target tone might pose challenges due to the onset response triggered by the preceding maskers. For instance, previous studies by Epp et al. [2013], Egger et al. [2019], and Kim and Epp [2023] used a preceding masker interval longer than 500*ms* to prevent an overlap between the onset response to the preceding noise and the response to the target tone.

Together with the interaction of preceding stream formation with CMR (e.g., *CMR*_*CC*_ vs *CMR*_*FC*_), we argue that the contribution of neural processes at each stage along the auditory pathway can differ depending on the properties of the auditory cues for stream formation. In addition, unlike our assumption that the changes in behavioral measures in long time scale can be linked to the high-level processing, the low-level processing can also show rather longer time scale (> 200*ms*). Even though the direct physiological evidence is lacking in this study, low-level processing has a potential in complex neural processing considering that various types of neurons exist in the CN, and these neurons are connected to each other forming neural circuits such as feed-forward excitation/inhibition, feedback inhibition, and mutual excitation [Oertel et al., 2011, Manis and Campagnola, 2018, Ngodup et al., 2020]. For instance, in the visual system, the adaptation to light contrast occurs at the retinal level (equivalent to the cochlea), ranging from 0.1 to 17 seconds [Baccus and Meister, 2002]. These time constants can be realized by different retinal cells such as bipolar cells, amacrine cells, and ganglion cells [Kohn, 2007]. In addition, it might worth it to note that to define “stream formation at high-level processing”, one can consider whether it results in perceptual stream or not. Furthermore, for the non-linear summation of CMR and BMLD, a possible neural basis can be a contralateral projection of a wideband inhibitor cell [Verhey et al., 2003, Ernst and Verhey, 2006, Ingham et al., 2006].

## 5. Conclusion

The present study investigated the effect of the preceding masker on CMR and BMLD by varying the length of the preceding masker. The effect of the preceding masker on the following sound has been suggested to be either the result of an adaptation effect at peripheral level (on short time scales) or system level (on long time scales). From the data acquired in this study, we showed that the time scale of the effect of preceding masker can range up to 500 ms on CMR, depending on the consistency of the comodulation cue in the preceding masker interval and the masked signal interval. However, the preceding stream formation did not affect BMLD induced by IPD of the target tone. This may indicate that the impact of the preceding stream formation may stem from neural processing at a peripheral level or from interactions between subcortical processes (e.g., the CN) and cortical neural processes at the A1 level, rather than solely from higher-order processing at the systemic level (e.g., temporal integration).

## Acknowledgments

This study was supported by a scholarship from the Technical University of Denmark to HK.

## 6. Supplement

